# Genome-scale metabolic models reveal phylogenetic conservation of amino acid fermentation-linked metabolic traits in soil bacteria

**DOI:** 10.64898/2026.11.02.763679

**Authors:** Young C. Song, Christian Ayala-Ortiz, Kelly G. Stratton, Arjun Chakrawal, Emily B. Graham

**Affiliations:** Integrated Discovery Sciences Directorate, Pacific Northwest National Laboratory, Richland, WA 99354, United States; School of Biological Sciences, Washington State University, Richland, WA, USA

## Abstract

Soil microorganisms drive biogeochemical cycling of carbon, nitrogen, and other elements, but the partitioning of metabolic fluxes among taxa remains poorly understood, limiting our ability to link community composition to ecosystem function. Genome-scale metabolic models (GEMs) offer a mechanistic framework to connect taxa and their metabolic pathways to these fluxes. However, heterogeneity of soil communities has hampered recovery of high-quality metagenome-assembled genomes (MAGs), constraining application of flux balance analysis (FBA) to soil metagenomes and skewing available GEMs towards cultured model organisms. This gap limits our understanding of soil microbial physiology and responses to disturbance. Here, we present a collection of GEMs for 550 MAGs (497 bacterial, 53 archaeal) recovered from 94 soil samples across the continental United States. As an example application, we used FBA to estimate uptake and secretion of 44 commonly exchanged metabolites and tested whether these fluxes are phylogenetically conserved. While the overall bacterial phylogenetic signal was weak (median r = 0.029), eight metabolites showed significant conservation: amino acids (L-methionine, L-leucine, L-isoleucine), fermentation intermediates (pyruvate, acetaldehyde, ethanol), and amino acid degradation by-products NH_3_ and CO_2_. Most bacterial MAGs were predicted to import branched-chain amino acids, and a subset also secreted pyruvate and ethanol, identifying candidate cross-feeding interactions. Thus, fermentation-linked traits may be phylogenetically conserved and predictable from community composition, whereas most other metabolites showed no detectable signal. Ultimately, this resource enables genome-resolved prediction of soil microbial metabolisms, from cross-feeding and carbon-use efficiency to community responses to disturbance, providing a mechanistic bridge between microbial taxonomy and ecosystem-scale biogeochemistry.

## Main

Soil microbial communities are highly diverse, and the vast majority of their members remain uncultured, presenting a bottleneck in understanding their metabolic capabilities, interactions, and responses to environmental change [1,2]. While metagenomics has greatly expanded the catalogue of soil microbial genomes through the recovery of metagenome-assembled genomes (MAGs), translating gene content into quantitative predictions of metabolic function remains a major challenge. A comprehensive set of genome-scale metabolic models (GEMs) for soil microbes would enable the cross-feeding, niche partitioning, and disturbance responses of these uncultured members to be explored *in silico* [3]. Metabolic modelling with GEMs has yielded mechanistic insights in other microbial systems, revealing conserved amino acid and B-vitamin cross-feeding among ocean bacterioplankton [3] and diet-dependent short-chain fatty acid production in the human gut microbiome [4]. Comparable analyses in soils, however, have been limited by the absence of an equivalent resource, as existing GEMs largely represent cultured organisms rather than native soil taxa. Here, we address this gap by presenting GEMs for 550 MAGs recovered from 94 soil samples across the continental United States, providing a resource for generating and testing mechanistic hypotheses about soil microbial metabolism.

Deciphering the relationship between phylogeny and trait distribution is a central question in microbial ecology, as it could enable prediction of how changes in community composition affect ecosystem function. In microorganisms, gene loss, convergent evolution, and horizontal gene transfer can produce polyphyletic trait distributions, yet these processes also leave clade-specific genomic signatures, such as reduced genome size, variable G+C content, and conserved pathway configurations [5]. Indeed, the depth of conservation scales with trait complexity. Complex traits such as photosynthesis and methanogenesis are confined to a few deep clades, whereas the ability to use simple carbon substrates is highly dispersed [5,6]. These trait-based analyses, however, have relied on the presence or absence of genes or phenotypes, which may not capture how these capabilities are expressed through an organism’s metabolic network. GEMs coupled with flux balance analysis (FBA), which predicts reaction fluxes from network stoichiometry under defined constraints, offer a way to evaluate phylogenetic conservation at the level of quantitative metabolic function.

To build a soil metabolic modelling resource and test the phylogenetic conservation of metabolite fluxes, we reconstructed a GEM for 550 of 551 dereplicated MAGs generated from 94 soil samples collected across the continental United States (**Table S1**) by mapping annotated genes to biochemical reactions, and used FBA to estimate steady-state reaction fluxes. Simulations were run in KBase [2,7] under default complete media, yielding uptake and secretion profiles for 255 metabolites that reflect theoretical metabolic potential, independent of variation in soil chemistry and environmental conditions. The resulting GEM collection spans 497 bacterial MAGs from 23 phyla, dominated by *Actinomycetota* (n = 128), *Pseudomonadota* (n = 108), and *Acidobacteriota* (n = 69), and 53 archaeal MAGs, nearly all (n = 49) within the ammonia-oxidizing order *Nitrososphaerales* (**Table S2**). A maximum-likelihood phylogeny of the bacterial MAGs was built including up to three high-quality GTDB reference genomes [8] per genus with three or more MAGs, where available (n = 88 genomes from 31 genera; **Figure 1**). The most abundant orders were *Rhizobiales* (n = 59), *Chthoniobacterales* (n = 39), and the candidate order UBA4738 (n = 22) (**Table S2**). Together, these GEMs provide a foundation for characterizing uncultured soil microorganisms at a continental scale and testing how microbial interactions and environmental responses shape soil biogeochemistry.

**Figure 1.**
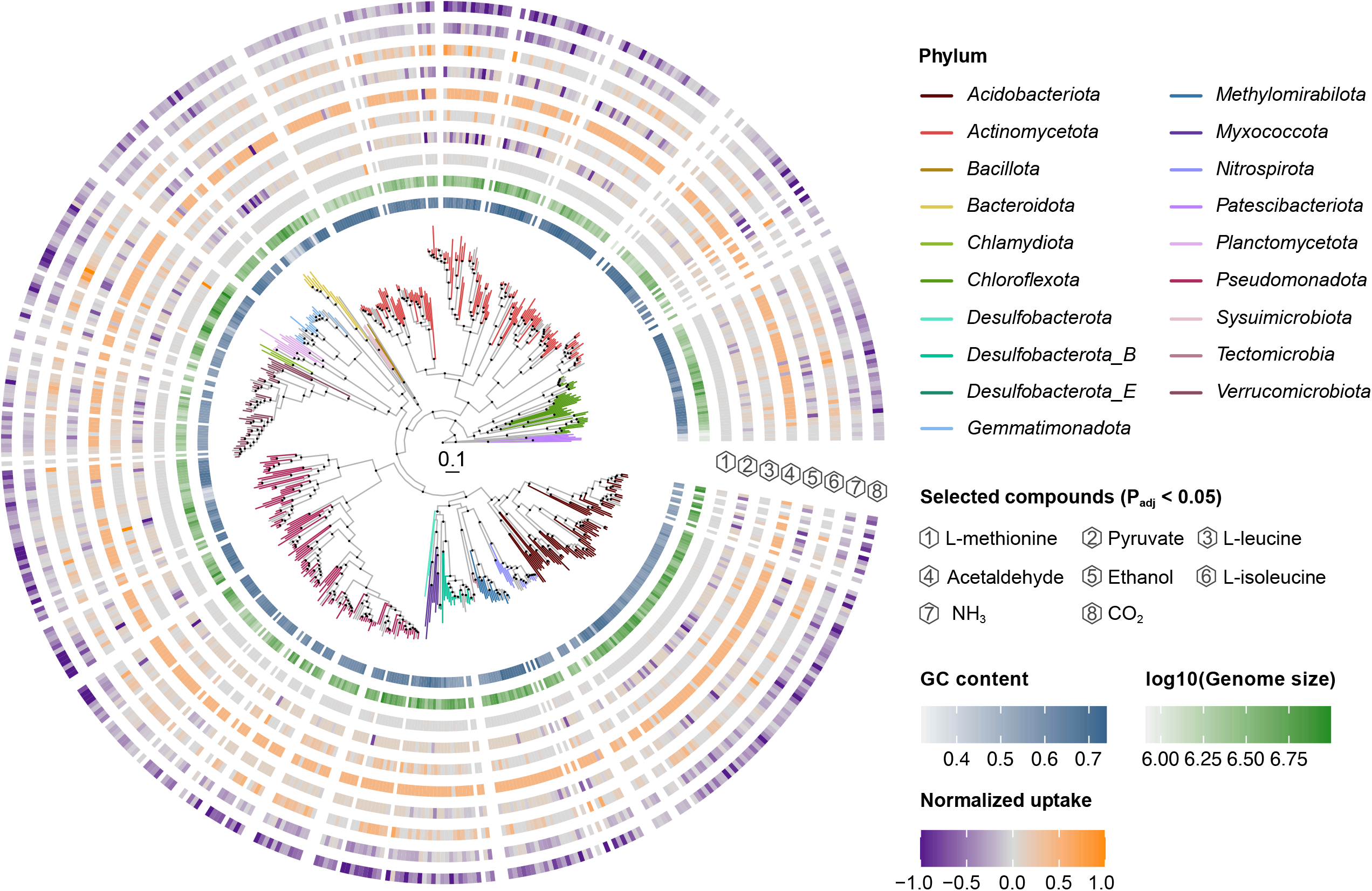
Maximum-likelihood phylogeny of 498 bacterial metagenome-assembled genomes (MAGs) and 88 GTDB species-representative reference genomes (up to three per genus represented by three or more MAGs, where available), inferred with IQ-TREE from the bac120 marker-gene alignment and rooted on *Patescibacteriota*. Branches are coloured by phylum for the 19 phyla represented by two or more MAGs; phyla represented by a single MAG are shown in grey. Internal nodes with ultrafast bootstrap support ≥ 90% are marked with black dots, and the scale bar indicates substitutions per site. The two innermost rings show GC content (0–1 scale) and log_10_-transformed genome size of each MAG, each on its own continuous colour scale. The eight outer rings show each MAG’s predicted exchange flux for one metabolite, coloured on a diverging scale (purple, net secretion; grey, no net exchange; orange, net uptake) after symmetric min–max normalization, in which each metabolite’s fluxes are divided by their maximum absolute value to give a signed range of −1 to 1. These are the eight metabolites with significant phylogenetic signal (Benjamini–Hochberg-adjusted P < 0.05; Spearman Mantel test between patristic distance and normalized flux difference, 999 permutations), ordered from innermost to outermost by decreasing whole-tree Mantel r, from L-methionine (r = 0.105) to CO_2_ (r = 0.041). Reference genomes, and one MAG for which no functional model could be generated, have no FBA predictions and appear as gaps in these rings.

Comparative genomic analyses have shown that predicted phenotypes, including growth, carbon-use efficiency, and substrate preference, are partly structured by phylogeny [9–11]. However, these studies relied largely on cultured reference genomes and aggregate or binary phenotypes, leaving questions regarding the phylogenetic conservation of metabolite-level exchange in uncultured soil taxa unresolved. We tested whether predicted fluxes of 44 commonly exchanged metabolites track phylogenetic distance, using Mantel tests between pairwise patristic distances and pairwise differences in normalized flux (**Supplementary Methods**; **Table S3**). Archaeal MAGs showed no significant associations (minimum P_adj_ = 0.861), likely reflecting their low number and limited diversity, so subsequent analyses focused on bacteria. Most metabolite fluxes showed no significant phylogenetic signal (median r = 0.029). However, eight metabolites showed significantly more similar fluxes among closely related MAGs (P_adj_ < 0.05): L-methionine (r = 0.105), pyruvate (r = 0.102), L-leucine (r = 0.075), acetaldehyde (r = 0.074), ethanol (r = 0.062), L-isoleucine (r = 0.052), NH_3_ (r = 0.044), and CO_2_ (r = 0.041) (**Figure 2, Figure S1**). These comprise amino acids, fermentation intermediates, and the amino acid degradation by-products NH_3_ and CO_2_, suggesting that traits linked to amino acid fermentation may be phylogenetically conserved.

**Figure 2.**
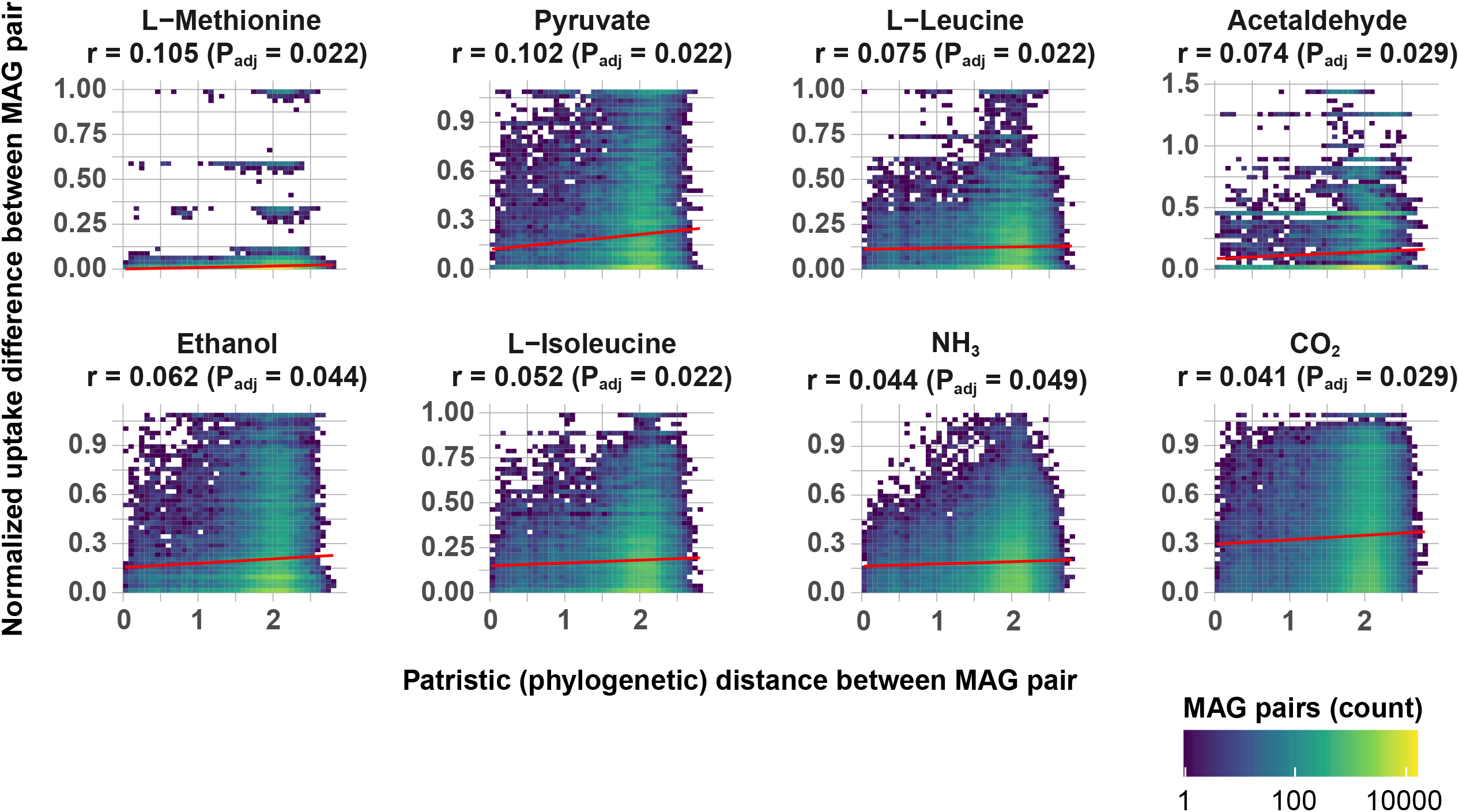
Phylogenetic distance versus difference in predicted metabolite exchange for the eight metabolites with significant phylogenetic signal (Benjamini–Hochberg-adjusted P < 0.05). Each panel shows one metabolite. The horizontal axis is the patristic distance between each pair of bacterial MAGs, and the vertical axis is the absolute difference in normalized flux (uptake or secretion, scaled to −1 to 1) between the same pair. MAG pairs are binned (40 × 40 bins), with colour indicating the number of pairs per bin on a log_10_ scale. The red line shows an ordinary least-squares regression fitted to all MAG pairs, with its 95% confidence band, for visual reference; significance was assessed by Mantel tests (Spearman correlation, 999 permutations), and each panel is labelled with the corresponding Mantel r and P_adj_. Panels are ordered by decreasing Mantel r.

Patterns among the significant metabolites further point to the import of branched-chain amino acids (BCAAs) coupled with the excretion of fermentation products (**Figure 1**). Uptake of L-leucine and/or L-isoleucine was predicted for 70% of bacterial MAGs (n = 348/497), whereas no MAG secreted either; uptake was most prevalent in *Verrucomicrobiota* (n = 40/42), followed by *Desulfobacterota_B* (n = 11/15), *Chloroflexota* (n = 27/38), and *Actinomycetota* (n = 90/128). Concurrent secretion of pyruvate and ethanol was predicted for 66 MAGs (13%), most frequently within *Verrucomicrobiota* (n = 12/42), *Gemmatimonadota* (n = 5/17), *Acidobacteriota* (n = 14/69), and *Actinomycetota* (n = 21/128). Importing BCAAs would relieve the anabolic demand that their biosynthesis places on pyruvate and acetyl-CoA [12], potentially freeing surplus pyruvate for excretion. Acetaldehyde uptake, predicted for 90% of bacterial MAGs, may further supply acetyl-CoA independently of pyruvate via acylating aldehyde dehydrogenase [13]. MAGs combining BCAA import with pyruvate and ethanol secretion (n = 36) spanned 54–73% GC (**Figure 1**) and did not differ in GC content from other bacterial MAGs (Mann–Whitney U test, P = 0.33), suggesting that this pattern is decoupled from the GC signatures associated with substrate preference in other bacteria [9]. These predictions thus outline a coupled BCAA-fermentation exchange strategy shared across eight bacterial phyla.

In summary, we present GEMs for 550 soil MAGs, providing a resource for exploring mechanistic hypotheses about soil microbial metabolism. Despite the overall weak phylogenetic signal in predicted metabolite exchange, traits linked to amino acid fermentation showed significant phylogenetic conservation across the bacterial tree. Together, these results identify candidate cross-feeding interactions for community-level modelling and suggest that soil bacterial community composition may be informative for predicting specific metabolic functions, offering a phylogenetically resolved framework for modelling soil biogeochemistry under changing environmental conditions.

## Supporting information

graphical_abstract

## Acknowledgements

Soil data was provided by the Molecular Observation Network (MONet) at the EMSL (https://doi.org/10.46936/10.25585/60008970). MONet is conducted in collaboration with Joint Genome Institute (https://ror.org/04xm1d337), a DOE Office of Science user facility supported by the Office of Science of the U.S Department of Energy operated under Contract DE-AC02-05CH11231. The MONet database is an open, FAIR, and publicly available compilation of the molecular and microstructural properties of soil. Data in the MONet open science database can be found at https://sc-data.emsl.pnnl.gov. The National Ecological Observatory Network is a programme sponsored by the U.S. National Science Foundation and operated under cooperative agreement by Battelle. A subset of samples used in this research were obtained through NEON Research Support Services. The KBase narrative used for this study was implemented with assistance of Zach Crockett at the ORNL.

## Author Contributions

YCS, CA and EBG conceived the project and directed the main experimental work. YCS conducted the bioinformatics analyses, including phylogenetic inferences of the MAGs, the FBA and the Mantel correlation analyses. CA and KGS provided guidance during statistical analyses of the metagenomic data. YCS and EBG wrote the manuscript together, while CA, KGS and AC provided constructive input on the structure and scientific accuracy of the manuscript.

