## Supplementary Information for "Genome-scale metabolic models reveal phylogenetic conservation of amino acid fermentation-linked metabolic traits in soil bacteria"

**Supplementary Information for the article, “Genome-scale metabolic models reveal phylogenetic conservation of amino acid fermentation-linked metabolic traits in soil bacteria”**

Young C. Song[1], Christian Ayala-Ortiz[1], Kelly G. Stratton[1], Arjun Chakrawal[1], *Emily B. Graham[1,2]

1. Integrated Discovery Sciences Directorate, Pacific Northwest National Laboratory, Richland, WA 99354, United States.

2. School of Biological Sciences, Washington State University, Richland, WA, USA

### Supplementary Methods

#### Generation of metagenome-assembled genomes and their phylogenetic placements

We generated metagenome-assembled genomes (MAGs) using metaGEM [1] from 94 soil samples from 60 sites, representing 15 distinct ecoregions across the continental United States (**Table S1**) [2], with an average of 1.20 × $10^8$ raw reads per sample. Default parameters were used for each step of the prokaryotic MAG generation process, unless otherwise specified. Short read quality filtering and adapter trimming were applied to the set of reads using fastp (v0.20.0) [3]. The filtered reads were then assembled into contigs using MEGAHIT (v1.2.9) [4], with the -*-presets meta-sensitive* and --*min-contig-len 1000* flags. Contig coverage across samples was calculated by first indexing the contigs with the "bwa index" command, then cross-mapping the filtered reads to the contigs with bwa-mem (v0.7.17) [5]. We then used the "samtools view" command with the -*Sb* flag to convert the SAM files generated from the bwa tools to BAM format, which were subsequently sorted using the "samtools sort" command. The contig coverages were then generated from BAM files, and then together with the assemblies, were used as input for the binning tools, MetaBAT2 (v2.12.1) [6], MaxBin2 (v2.2.5) [7] and CONCOCT (v1.1.0) [8]. The best version of each bin from the output of these binning tools was extracted using the "bin_refinement" module of metaWRAP (v1.2.3) [9], with -*x 10* -*c 50* flags, yielding a set of bins with completeness and contamination thresholds of ≥ 50% and ≤ 10%, respectively. We then applied the "reassemble_bins" module to improve the quality of the bins selected from the refinement step. The 830 MAGs generated from these steps were dereplicated using galah (v0.4.0) [10] at 95% average nucleotide identity, a threshold commonly used to delineate species, yielding 551 clusters; the representative MAG for each cluster was selected based on CheckM2 [11] completeness and contamination. According to the Minimum Information about a Metagenome-Assembled Genome (MIMAG) standards [12], 82 (14.9%) of the representative MAGs were high quality and 469 (85.1%) were medium quality.

The 551 MAGs were taxonomically classified using the Genome Taxonomy Database Toolkit (GTDB-Tk, v2.5.2) [13] against GTDB release 10 (R10-RS226) [14], which assigned 498 MAGs to Bacteria and 53 to Archaea (**Table S2**). To place the MAGs in a broader phylogenetic context, reference genomes were selected from GTDB for each genus represented by three or more MAGs. For each such genus, up to three species-representative genomes meeting MIMAG high-quality criteria (≥ 90% completeness and < 5% contamination, as estimated by CheckM2) were chosen where available, prioritizing the highest completeness and lowest contamination. This yielded 88 bacterial reference genomes spanning 31 of the 39 eligible genera across 9 phyla, and 18 archaeal reference genomes spanning all 6 eligible genera (**Table S2**).

Maximum-likelihood phylogenies of the MAGs and reference genomes were inferred from the trimmed concatenated alignments of the 120 bacterial (bac120) and 53 archaeal (ar53) marker genes produced by GTDB-Tk, using IQ-TREE (v3.1.3) [15] with models selected by ModelFinder [16] according to the Bayesian information criterion (LG+F+R10 for Bacteria and Q.INSECT+F+R6 for Archaea) and 1,000 ultrafast bootstrap replicates. The bacterial tree was rooted using the phylum *Patescibacteriota* as the outgroup [17].

**Construction of genome-scale metabolic models using the MAGs**

To construct metabolic models for the 551 MAGs, we built a metabolic prediction narrative using applications available in the DOE Systems Biology Knowledgebase (KBase) [18]. Within this narrative, the MAGs were annotated using RAST-Tk (Rapid Annotation using Subsystems Technology-Toolkit, v1.73) [19], generating annotation objects that were used as input for the OMEGGA model-building application, which reconstructs draft models from ModelSEED templates [20]. For constructing all models, gap filling was applied to add the minimal set of reactions required for biomass production. The resulting models comprise matrices of metabolic reactions, their associated biochemical compounds, and gene-protein-reaction associations, based on the genes identified in the RAST-Tk step and the gapfilling process. Finally, the flux-balance-analysis (FBA) [21] application (fba_tools) was used to estimate steady-state reaction fluxes for each model by maximizing biomass production (bio1) under the KBase default complete medium, in which every compound the model can transport is available (uptake bound of 100 and secretion bound of 1,000 mmol $gDW^{-1}$ $h^{-1}$). Flux variability analysis was enabled and flux minimization was applied. The FBA output includes the predicted growth rate (biomass flux), reaction fluxes, and uptake and secretion fluxes of extracellular metabolites.

**Phylogenetic signal in predicted metabolite exchange**

To test whether predicted metabolite exchange is phylogenetically conserved, exchange fluxes from the FBA simulations were compiled across MAGs separately for Bacteria and Archaea. For one bacterial MAG (PPRH-TOP_bin.5.o), a draft model was reconstructed using the Gram-negative template, but gap filling did not yield a solution supporting biomass production, and FBA predicted no growth; this MAG was therefore excluded, leaving 497 bacterial and 53 archaeal MAGs. Metabolites without a predicted flux in one-third or more of the MAGs were removed, and remaining missing values were set to zero. This retained 50 of 254 metabolites for Bacteria and 67 of 141 for Archaea. Because FBA fluxes are signed (positive values indicate uptake and negative values indicate secretion) and many reach the uptake bound, each metabolite's fluxes were scaled by their maximum absolute value. This symmetric scaling gives values from −1 to 1 and preserves both sign and true zeros. Metabolites with zero variance across MAGs could not be tested and were excluded: six for Bacteria ($Cd^{2+}$, $Hg^{2+}$, $Na^{+}$, $Ni^{2+}$, Pb, and glycogen) and nine for Archaea. This left 44 bacterial and 58 archaeal metabolites.

Pairwise patristic distances between MAGs were calculated with *ape* (v5.8.1) [22] from the ultrafast bootstrap consensus trees of the bac120 and ar53 alignments, read with *treeio* (v1.32.0) [23], after reference genome tips were pruned so that only MAGs with FBA predictions were compared. For each metabolite, a pairwise dissimilarity matrix was built from the absolute differences in normalized flux between MAGs. Mantel tests between the phylogenetic and flux-dissimilarity matrices were performed with *vegan* (v2.7.5) [24], using Spearman rank correlation because both matrices are non-negative and skewed, with 999 permutations (random seed = 1). A positive Mantel r indicates that more closely related MAGs have more similar exchange of that metabolite. P-values were corrected for multiple testing across metabolites within each domain using the Benjamini–Hochberg false discovery rate [25], and metabolites with $P_{adj} < 0.05$ were considered significantly conserved (**Table S3**). Eight trace ions ($Mg^{2+}$, $Co^{2+}$, $K^{+}$, $Mn^{2+}$, $Cl^{-}$, $Ca^{2+}$, $Cu^{2+}$, and $Zn^{2+}$) with identical flux profiles across MAGs, and the enterobactin/Fe-enterochelin pair, produced identical dissimilarity matrices. All were retained as separate tests, which makes the correction conservative.

**Genomic GC content of metabolite-exchange patterns**

To assess whether the combined import of the branched-chain amino acids (BCAAs) L-leucine and/or L-isoleucine and secretion of fermentation products was associated with genomic GC content, bacterial MAGs were classified by their predicted fluxes in the filtered FBA dataset. MAGs were assigned to this pattern if they had positive fluxes (uptake) for L-leucine and/or L-isoleucine together with negative fluxes (secretion) for both pyruvate and ethanol ($n = 36$ of 497). GC content, estimated by CheckM2, was compared between these and all other bacterial MAGs ($n = 461$) using a two-sided Wilcoxon rank-sum (Mann–Whitney U) test in R [26].

### Data Availability

Sequences of the 551 dereplicated MAGs and their genome-scale metabolic models (550 MAGs), including model reactions and compounds, FBA-predicted reaction fluxes, and compound uptake and secretion profiles, are available on Zenodo (https://doi.org/10.5281/zenodo.23480528). The R code and input data used for the phylogenetic analyses, figures, and supplementary tables are available on GitHub (https://github.com/EMSL-MONet/The_1000_Soils_MAG_GEM_Phylo_Signal). The KBase narrative developed to generate GEMs and conduct FBA analyses can be accessed at https://doi.org/10.25982/3020148.

**Supplementary Tables**

All tables are provided in a separate file, Supplementary_Data.xlsx.

**Table S1.** Locations of soil samples, from which MAGs were recovered. Information in the table includes site codes and names ('Site_Code' and 'Site_Name'), NEON ecoregion ('NEON_ecoregion'), geographical coordinates (latitude and longitude), and elevation in metre (‘Elev.m’).

**Table S2**. Taxonomy and genome quality of the dereplicated MAGs and phylogenetic reference genomes. GTDB-Tk (v2.5.2; GTDB R10-RS226) classifications and CheckM2 quality metrics for the 551 dereplicated MAGs (498 bacterial and 53 archaeal), including completeness, contamination, genome size, GC content, assembly statistics, and MIMAG quality tier (82 high-quality and 469 medium-quality). Also listed are the GTDB reference genomes included in the bacterial (n = 88, 31 genera) and archaeal (n = 18, 6 genera) phylogenies, with their GTDB taxonomy to genus level and CheckM2 completeness and contamination. Reference genomes were GTDB species representatives selected for each genus represented by three or more MAGs (up to three per genus where available; ≥ 90% completeness and < 5% contamination).

**Table S3**. Phylogenetic signal in predicted metabolite exchange across bacterial MAGs. Whole-tree Mantel correlations between pairwise patristic distance and pairwise difference in normalized FBA-predicted flux for 44 metabolites across 497 bacterial MAGs. Fluxes were scaled by each metabolite's maximum absolute value before computing pairwise differences. Mantel tests used Spearman rank correlation with 999 permutations, and p-values were adjusted across metabolites using the Benjamini–Hochberg false discovery rate ($P_{adj}$). Positive r indicates that more closely related MAGs exchange a metabolite more similarly. Metabolites are ordered by decreasing r; those with $P_{adj} < 0.05$ are shown in **Figure 2**, and the remainder in **Figure S1**. Six metabolites with zero variance across MAGs ($Cd^{2+}$, $Hg^{2+}$, $Na^{+}$, $Ni^{2+}$, Pb, and glycogen) could not be tested and are not listed.

**Supplementary Figures**

**Figure S1. Phylogenetic distance versus difference in predicted metabolite exchange for the remaining 36 tested bacterial metabolites (Benjamini–Hochberg-adjusted P ≥ 0.05).** Panels are constructed as in Figure 2 and ordered by decreasing Mantel r.
