## Supplementary material for "Genome-scale metabolic models reveal phylogenetic conservation of amino acid fermentation-linked metabolic traits in soil bacteria": FigS1

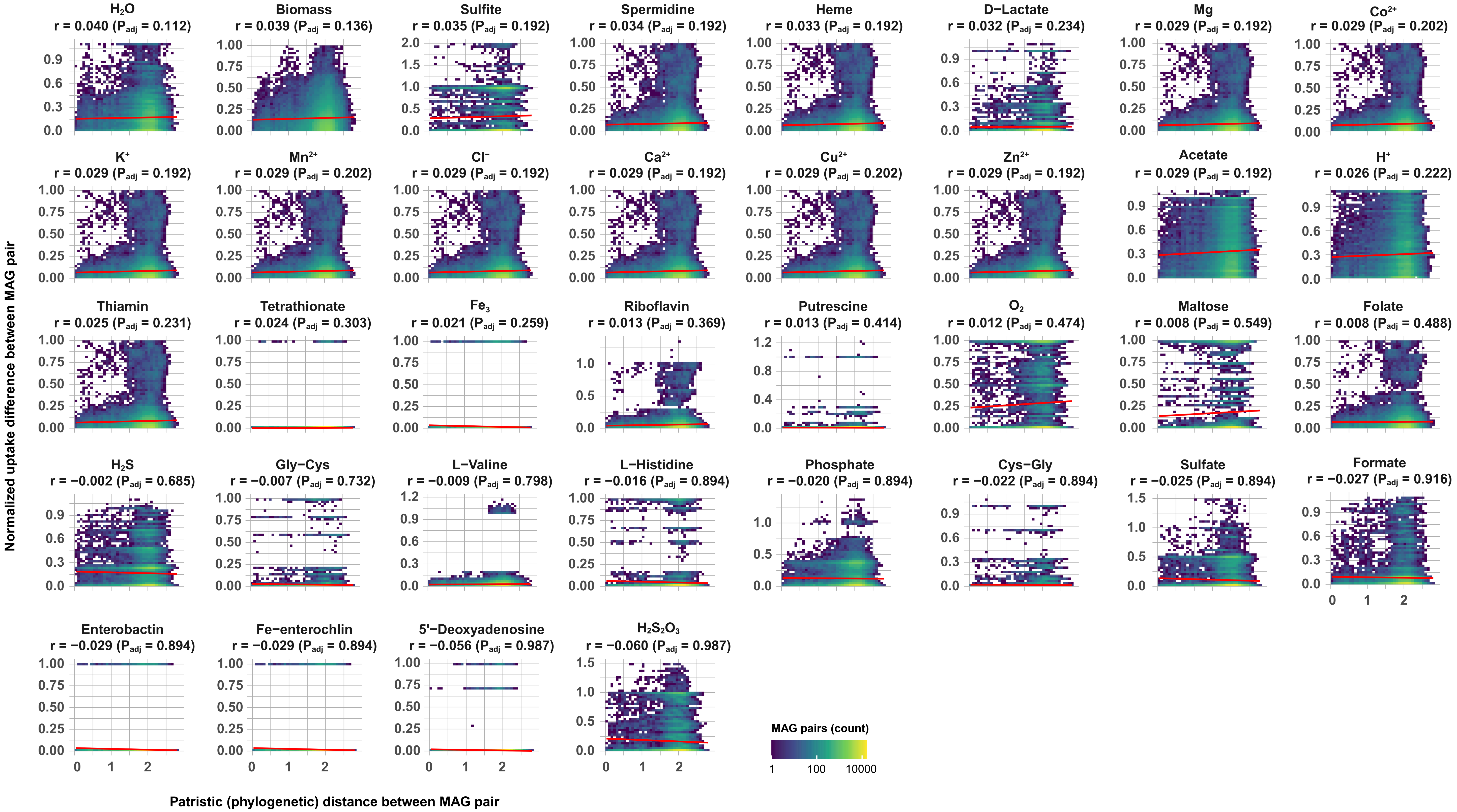

H2O r = 0.040 (Padj = 0.112)
Biomass r = 0.039 (Padj = 0.136)
Sulfite r = 0.035 (Padj = 0.192)
Spermidine r = 0.034 (Padj = 0.192)
Heme r = 0.033 (Padj = 0.192)
D−Lactate r = 0.032 (Padj = 0.234)
Mg r = 0.029 (Padj = 0.192)
Co2+ r = 0.029 (Padj = 0.202)
K+ r = 0.029 (Padj = 0.192)
Mn2+ r = 0.029 (Padj = 0.202)
Cl− r = 0.029 (Padj = 0.192)
Ca2+ r = 0.029 (Padj = 0.192)
Cu2+ r = 0.029 (Padj = 0.202)
Zn2+ r = 0.029 (Padj = 0.192)
Acetate r = 0.029 (Padj = 0.192)
H+ r = 0.026 (Padj = 0.222)
Thiamin r = 0.025 (Padj = 0.231)
Tetrathionate r = 0.024 (Padj = 0.303)
Fe3 r = 0.021 (Padj = 0.259)
Riboflavin r = 0.013 (Padj = 0.369)
Putrescine r = 0.013 (Padj = 0.414)
O2 r = 0.012 (Padj = 0.474)
Maltose r = 0.008 (Padj = 0.549)
Folate r = 0.008 (Padj = 0.488)
H2S r = −0.002 (Padj = 0.685)
Gly−Cys r = −0.007 (Padj = 0.732)
L−Valine r = −0.009 (Padj = 0.798)
L−Histidine r = −0.016 (Padj = 0.894)
Phosphate r = −0.020 (Padj = 0.894)
Cys−Gly r = −0.022 (Padj = 0.894)
Sulfate r = −0.025 (Padj = 0.894)
Formate r = −0.027 (Padj = 0.916)
Enterobactin r = −0.029 (Padj = 0.894)
Fe−enterochlin r = −0.029 (Padj = 0.894)
5'−Deoxyadenosine r = −0.056 (Padj = 0.987)
H2S2O3 r = −0.060 (Padj = 0.987)
Normalized uptake difference between MAG pair
Patristic (phylogenetic) distance between MAG pair
MAG pairs (count)
1
100
10000
