## Supplementary figures and images for "Genome-scale metabolic models reveal phylogenetic conservation of amino acid fermentation-linked metabolic traits in soil bacteria"

### graphical_abstract

#### Genome-scale metabolic models for soil microbial communities

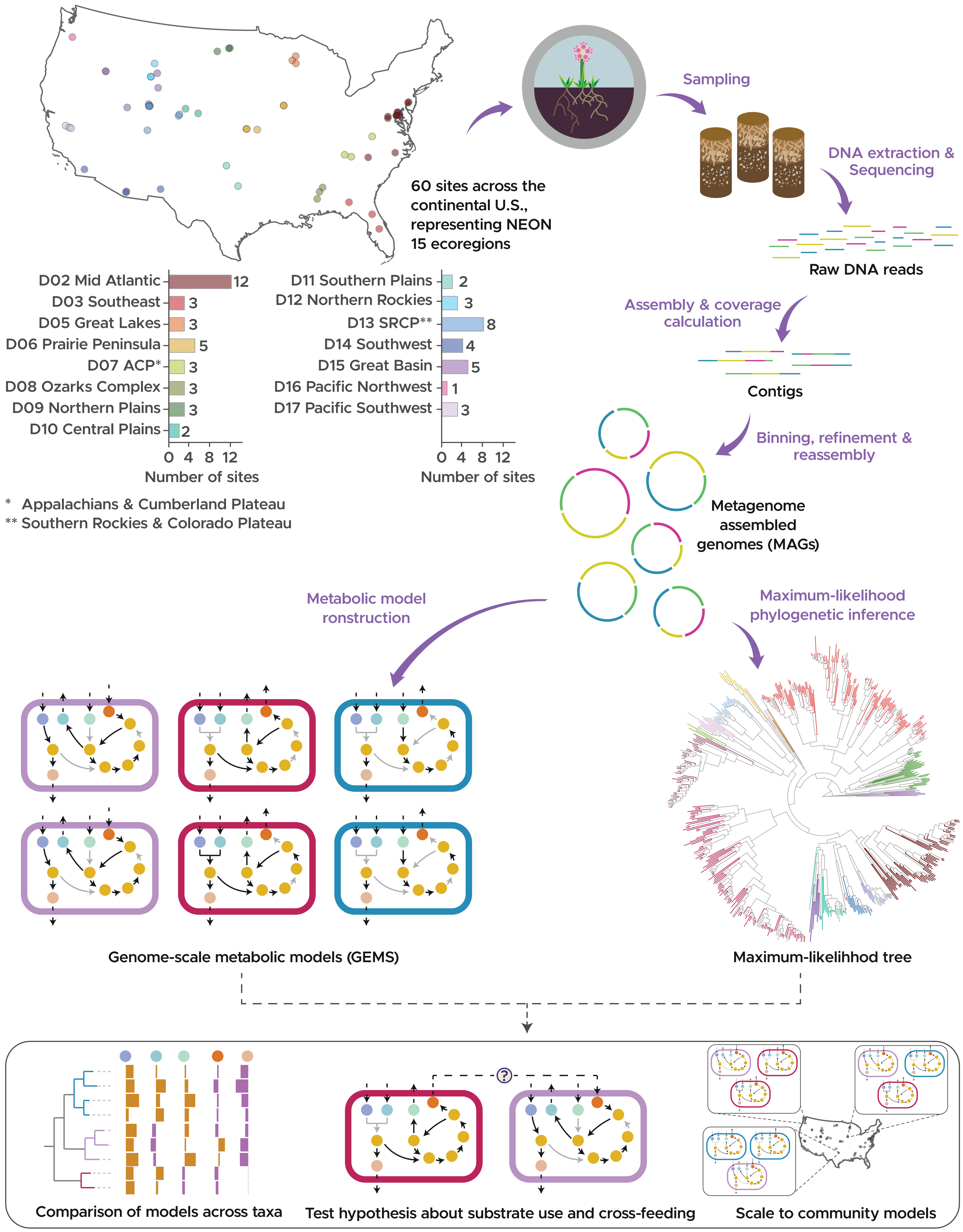
